# Optimizing Glycemic Control in Type 1 Diabetic Patients using a Deep Learning-Based Artificial Pancreas with a Secure Glucagon and Insulin Delivery System

**DOI:** 10.1101/2023.12.07.566476

**Authors:** Rohan Singh, Raj R. Rao

**Author notes:** **Contact information:** Rohan Singh Raj Rao.

## Abstract

Type 1 diabetes impacts millions worldwide, with some patients facing rapid fluctuations in their blood sugar levels. These fluctuations can negatively impact an individual’s quality of life and if untreated, can lead to nerve damage, coma, and death. While current methods have helped address hyperglycemia (high blood sugar), there has been less success with hypoglycemia (low blood sugar) and glucagon administration. To bridge this gap, an artificial pancreas with a novel insulin and glucagon pump was developed. Initiating the system, a personalized mobile app enables users to input meal carbohydrate and insulin bolus data. The data is then transmitted to a deep learning model that incorporates Continuous Glucose Monitor readings, along with carbohydrate and insulin data from the app. The two-layer Long Short-Term Memory network, developed in Python, accurately forecasts blood sugar levels on Ohio University’s OhioT1DM patient dataset and the UVa/Padova simulation’s data for a 30 minute-interval. An algorithm then utilizes the predictions to calculate optimal insulin and glucagon doses using metabolization formulas. To ensure system security, data is transmitted through a cloud-based MQ Telemetry Transport server and secured with industry-standard authentication and encryption methods. Finally, a microcontroller-based prototype accurately dispenses insulin and glucagon doses. The system kept in-silico patients at optimal levels for 38% longer and reduced dangerous levels by 22% compared to conventional controllers on an FDA-approved preclinical trial alternative simulation. By addressing both hypo and hyperglycemia, this real-time medical device can be a transformative tool for individuals with diabetes, enabling them to live healthier and more fulfilling lives.

## Background

Diabetes, one of the fastest-growing chronic diseases, affects over 537 million people worldwide and alters how the body metabolizes glucose (International Diabetes Federation, 2021). Normally, the pancreas produces insulin, a natural hormone that lowers blood sugar, after a meal to reduce blood sugar by allowing glucose to enter the body’s cells. However, type 1 diabetes (T1D) is a chronic condition characterized by the destruction of beta-cells in the pancreas which are responsible for insulin production (Roep, Bart O et al., 2021) (Figure 1). Because of this, patients with T1D may not produce enough insulin leading to rapid fluctuations in blood sugar levels which leads to the increased risk of hyperglycemia (high blood sugar) and hypoglycemia (low blood sugar). If left untreated, hyper and hypoglycemia can lead to severe symptoms such as nerve damage, coma, and death (Mathew, P., & Thoppil, D., 2022; Mouri MI & Badireddy M, 2023).

**Figure 1.**
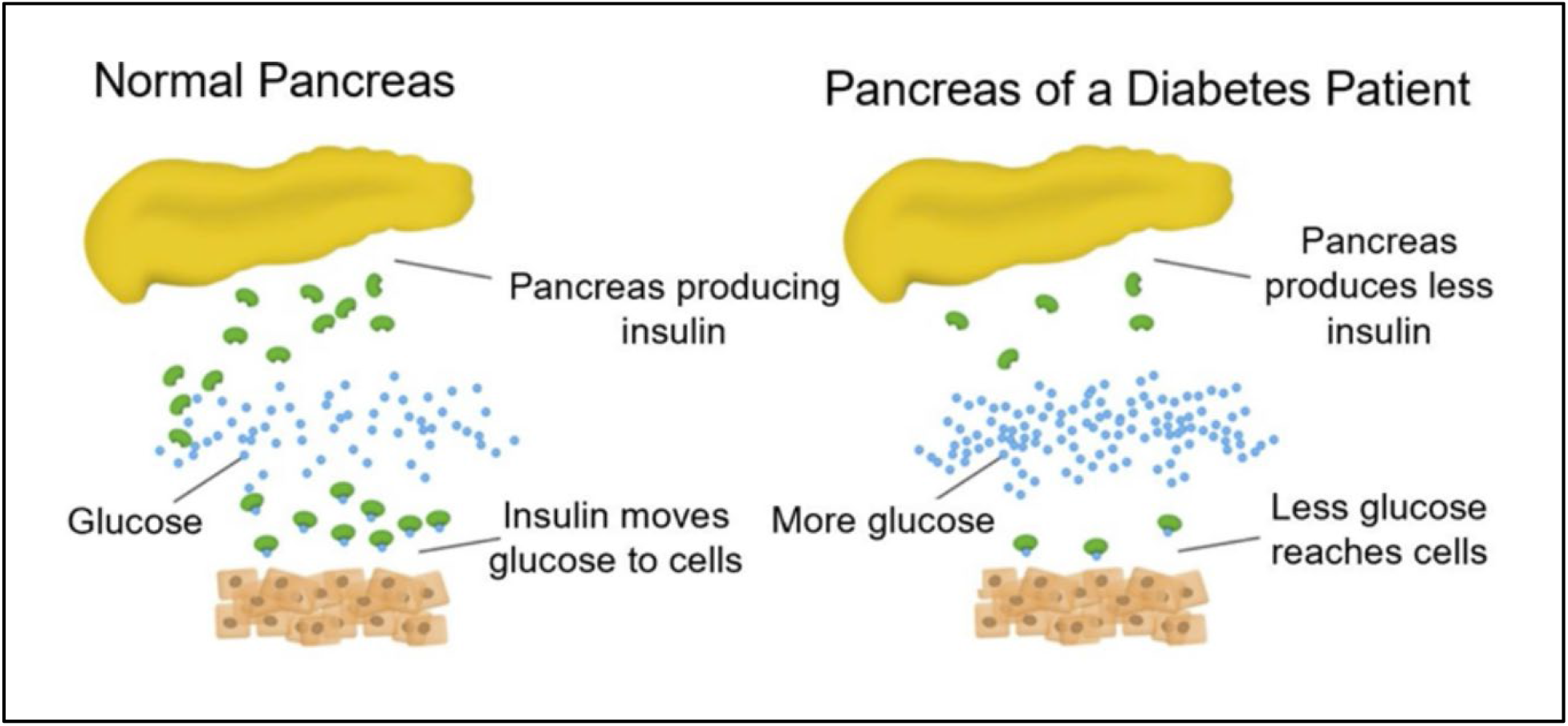
Poor glycemic control at the cellular and tissue level. Under normal conditions, a healthy pancreas produces adequate amounts of insulin that transport intracellular glucose to many cells and tissues such as the liver, muscle, and adipose tissue. In a diseased state, the ability of the pancreas to produce insulin is diminished, which results in a reduction in glucose to different cells and tissues, and an increase of glucose present in the bloodstream.

Furthermore, blood sugar fluctuations both physically and mentally affect patients and their caretakers. Patients with T1D must take regular insulin basal (longer-acting insulin) and bolus (rapid-acting insulin) doses after meals and at certain time intervals to maintain healthy blood sugar levels (JaneŽ, A., Guja, C., 2020). This process can be especially tedious during nighttime, when individuals with T1D and caretakers may need to wake up several times to address blood sugar fluctuations that can disrupt normal sleep patterns and reduce sleep quality which is associated with worse glycemic control (Chasens, E. R., 2013) (Figure 2). Adequate sleep is integral for maintaining optimal health and preventing disease progression. Poor glycemic control resulting from inadequate sleep can trap patients in a cycle of increased stress and even worse glycemic control (Sharma, S., & Kavuru, M., 2010; Farabi S. S., 2016) (Figure 3).

**Figure 2.**
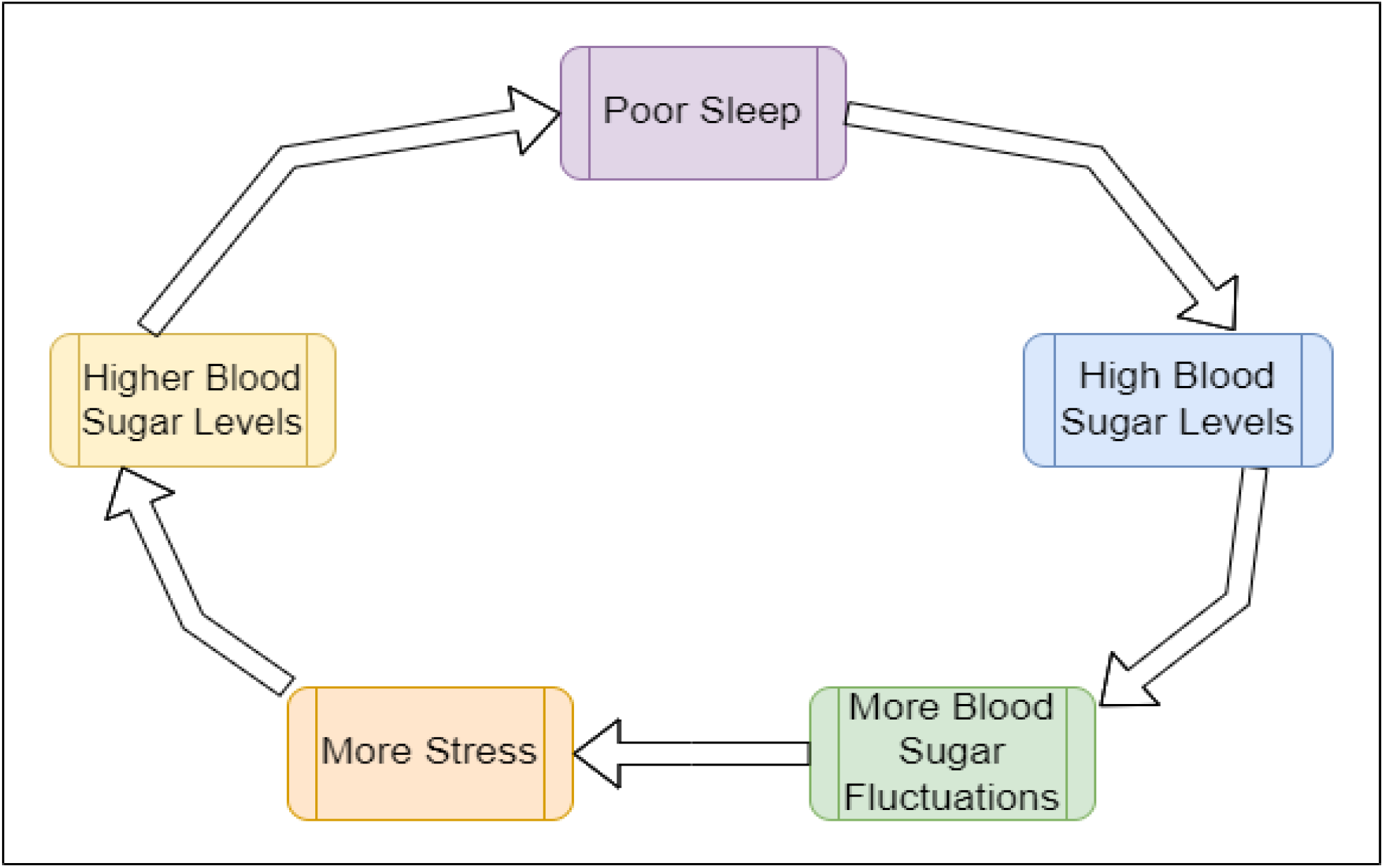
Cycle of poor glycemic control. In a T1D patient, poor sleep patterns contribute to high blood sugar levels, which in turn contribute to more blood sugar fluctuations. This then contributes to increased stress levels which can subsequently lead to higher blood sugar levels and poor sleep.

**Figure 3.**
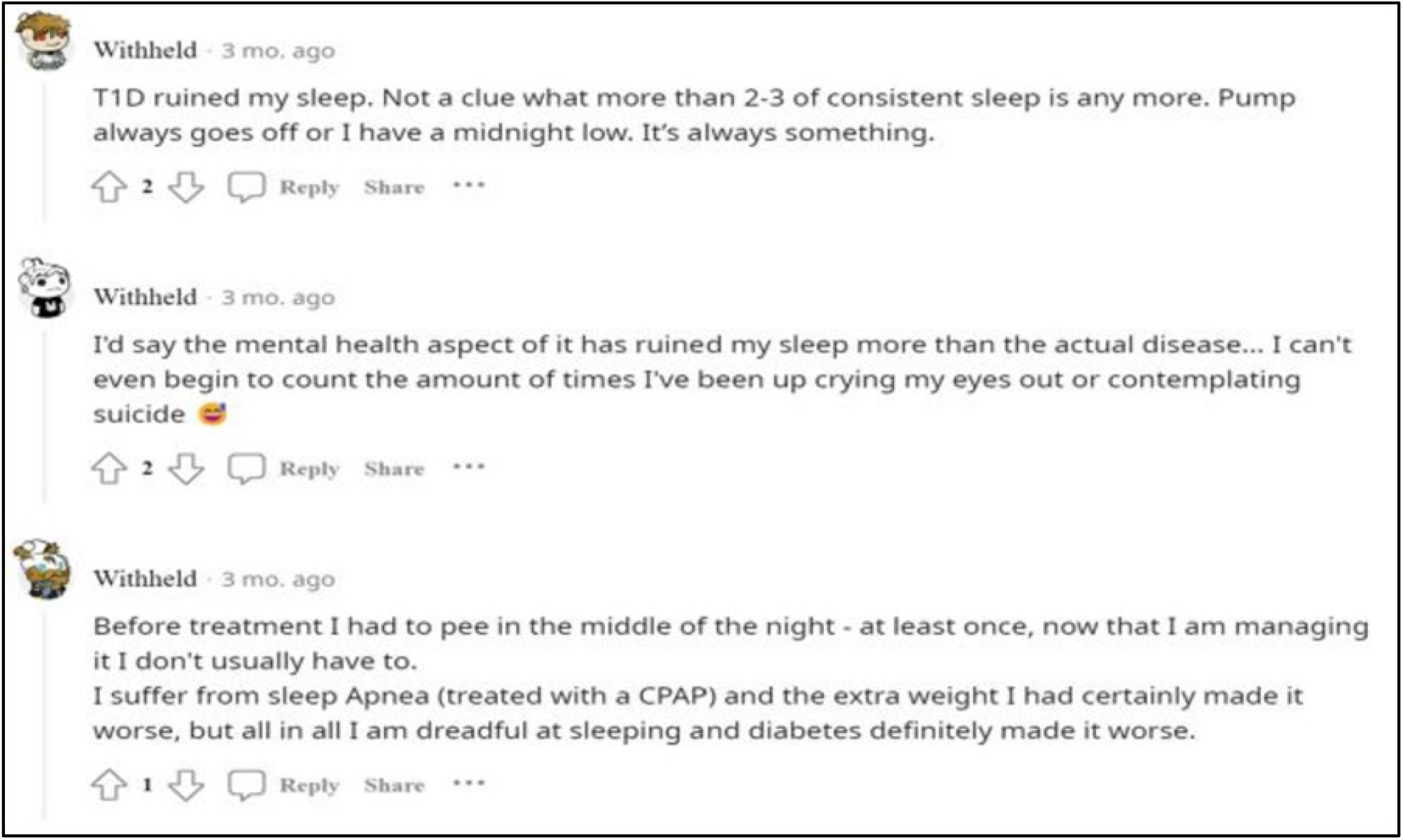
Responses from a Reddit post asking, “How has diabetes affected your sleep?” Anecdotal patient information indicates the potential for T1D to affect sleep patterns and potentially contribute to secondary complications because of T1D.

Many medical devices and associated technologies have been developed to aid patients in managing their diabetes. Continuous Glucose Monitors (CGMs) have been developed to provide users with a near real-time blood sugar reading, allowing users to view daily trends of their blood sugar. CGMs have been paired with mobile applications that present trend statistics to users in a simple, easy-to-understand manner (Miller, E. M., 2020). CGMs also opened new avenues of research by linking up with insulin pumps which are wearable, motorized devices that automatically deliver insulin doses subcutaneously through a small cannula. Insulin pumps utilize wireless network technology to be given dose amounts from a mobile application or external dosage calculation algorithm (Berget, C., et al., 2019).

Combined with a mobile application and an insulin pump, a CGM could detect an unhealthy blood sugar level, transmit that information through a wireless network to the insulin pump which in turn would administer the insulin dose, and then send that data over to the mobile application for the user and their healthcare provider to review. A system with a CGM, insulin pump, a network connecting all the components, and an optional mobile application is categorized as an artificial pancreas, or a closed-loop system (Figure 4) (Moon, S. J., 2021).

**Figure 4.**
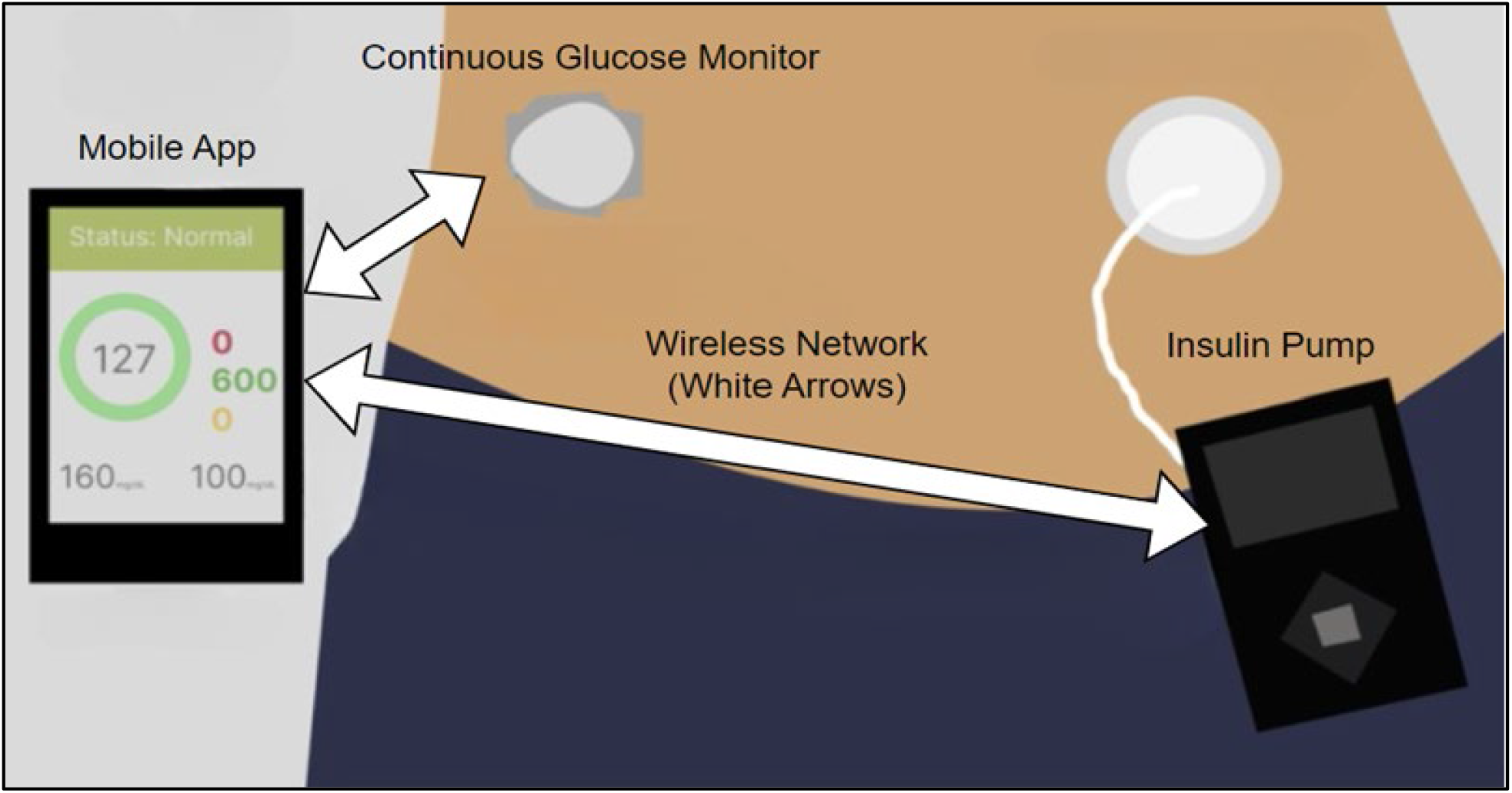
Components of an artificial pancreas or a closed-loop system. The continuous glucose monitor combines with an insulin pump to detect changes in glucose levels. These readings could then be transmitted through a wireless network to a mobile app that could allow the patient and healthcare provider to review them regularly (Figure adapted from Smith, C., 2023).

While artificial pancreas systems have been developed and implemented extensively, modern commercial systems have limitations. Firstly, current artificial pancreas systems only administer insulin and are therefore not designed to treat hypoglycemia effectively. In many cases, insulin is the preferred method for effectively managing hyperglycemia but not hypoglycemia (McDonnell, M. E., & Umpierrez, G. E. 2012). Because hypoglycemia is unaddressed in current systems, people using artificial pancreas systems often require manual intervention in cases of hypoglycemia. This is shown in recent novel artificial pancreas trials where despite advancements involving insulin dosage administration, groups of patients continued to experience moderate to severe hypoglycemia in each trial (Bionic Pancreas Research Group, Russell, S. J., 2022; Haidar A., 2019; Wadwa, R. P., et al., 2023). Currently, there are no clinically approved artificial pancreas systems utilizing both insulin and a different hormone or antidiabetic medication to fully address both hyper and hypoglycemia (Lindkvist, E. B., et al, 2023; Infante, M., et al, 2021) however, the hormone Glucagon has been identified as a possible method to address hypoglycemia. Glucagon is a hormone that gets secreted from the pancreatic alpha cells which contrasts with insulin by stimulating glucose production through increasing glucose concentration in the bloodstream, thus raising blood sugar levels. Glucagon is FDA-approved for treating hypoglycemia and is a popular choice for medical facilities and patients due to its simplicity and ease of use, being administered subcutaneously or intravenously (Morris, CH., and Baker, J., 2023). Dual hormone pumps, pump systems utilizing two hormones, have been tested and while the addition of hormones such as glucagon has shown promising results, hypoglycemia continues to be present in trial participants possibly due to imprecise treatment timings (Haidar, A., et al., 2015; Haider, A., 2019; Jessica, R, Castle., et al., 2018; Lindkvist, E. B., et al, 2023).

Blood sugar prediction algorithms, utilized by current artificial pancreas systems to better calculate future hormone doses, have difficulty predicting rapid fluctuations in blood sugar as these fluctuations are nonlinear and are influenced by numerous factors such as sleep quality, stress, insulin intake, and exercise (Adams O. P, 2013; Chasens, E. R., 2013; Surwit, R. S., et al., 1992). Improper blood sugar predictions can lead to inaccurate treatment, resulting in patients suffering from ineffective treatment. Machine learning has been utilized in various biomedical applications including disease detection and prediction using data from sensors, medical images, and biobanks, (Shah, P., et al., 2019; Battineni, G., et al., 2022). Deep learning is a subset of machine learning that shows great promise in extracting features and learning patterns from complex data (Cao, C., et al., 2018). Given the nonlinear nature of blood sugar, deep learning models could learn the blood sugar trends of each patient, allowing for more accurate and personalized predictions compared to current dosage algorithms.

A crucial but less frequently discussed problem is the threat of bad actors and hackers exploiting network vulnerabilities in medical devices such as artificial pancreas systems. Each component of an artificial pancreas system is in some way connected to a wireless network (Berget, C., et al., 2019). Within this network, physiological data, dose information, and sensitive user information are stored and transferred (O’Keeffe, D. T., 2015). Consequently, artificial pancreas systems could be vulnerable to cyber threats, potentially leading to the loss of sensitive patient data or system misuse in which a bad actor could gain control of the device’s functions and cause harm to the user (A. Tabasum., et al., 2018; Hassija, V., et al., 2021).

To address the limitations in modern-day artificial pancreas systems, a novel artificial pancreas that administers glucagon, in addition to insulin, to address both hypo and hyperglycemia while utilizing deep learning to accurately predict future blood sugar levels to calculate more accurate doses is proposed. A mobile application allows users to input everyday meal carbohydrate and insulin bolus amounts. This data is then sent to the deep learning model to predict future blood sugar levels. A dosage algorithm calculates the optimal insulin and glucagon doses using the model’s predictions. A cloud-based MQ Telemetry Transport (MQTT) server utilizes industry-standard encryption and authentication methods to connect the components of the system together while securing patient data and preventing unauthorized device access. Finally, a hardware insulin and glucagon pump administers the doses. The novel combination of these technologies allows for a more accurate, secure, and effective artificial pancreas system to allow individuals with T1D to maintain optimal glycemic levels for longer.

## Methodology

### Mobile Application

To start the system, a user-friendly mobile application was developed using the coding language Swift (Swift, 2023). The app has the capability to link with a CGM to display and record real-time blood sugar levels. Because carbohydrates from meals and insulin boluses have direct impacts on blood sugar levels by either raising or lowering levels respectively, the user can record the number of carbohydrates or the number of insulin boluses they took to allow the system to calculate optimal doses (Holesh, JE., et al., 2023; Donnor, T., et al., 2000). Users can specify their preferred blood sugar level range while also inputting personal factors such as insulin sensitivity, glucagon sensitivity, and correction factors, provided by the user’s healthcare providers, which are all taken into account by the dosage algorithm. Additional features such as calendars, reminders, and blood sugar visuals are also provided. If needed, patient data can be viewed and downloaded by the patient’s healthcare providers (Figure 5).

**Figure 5.**
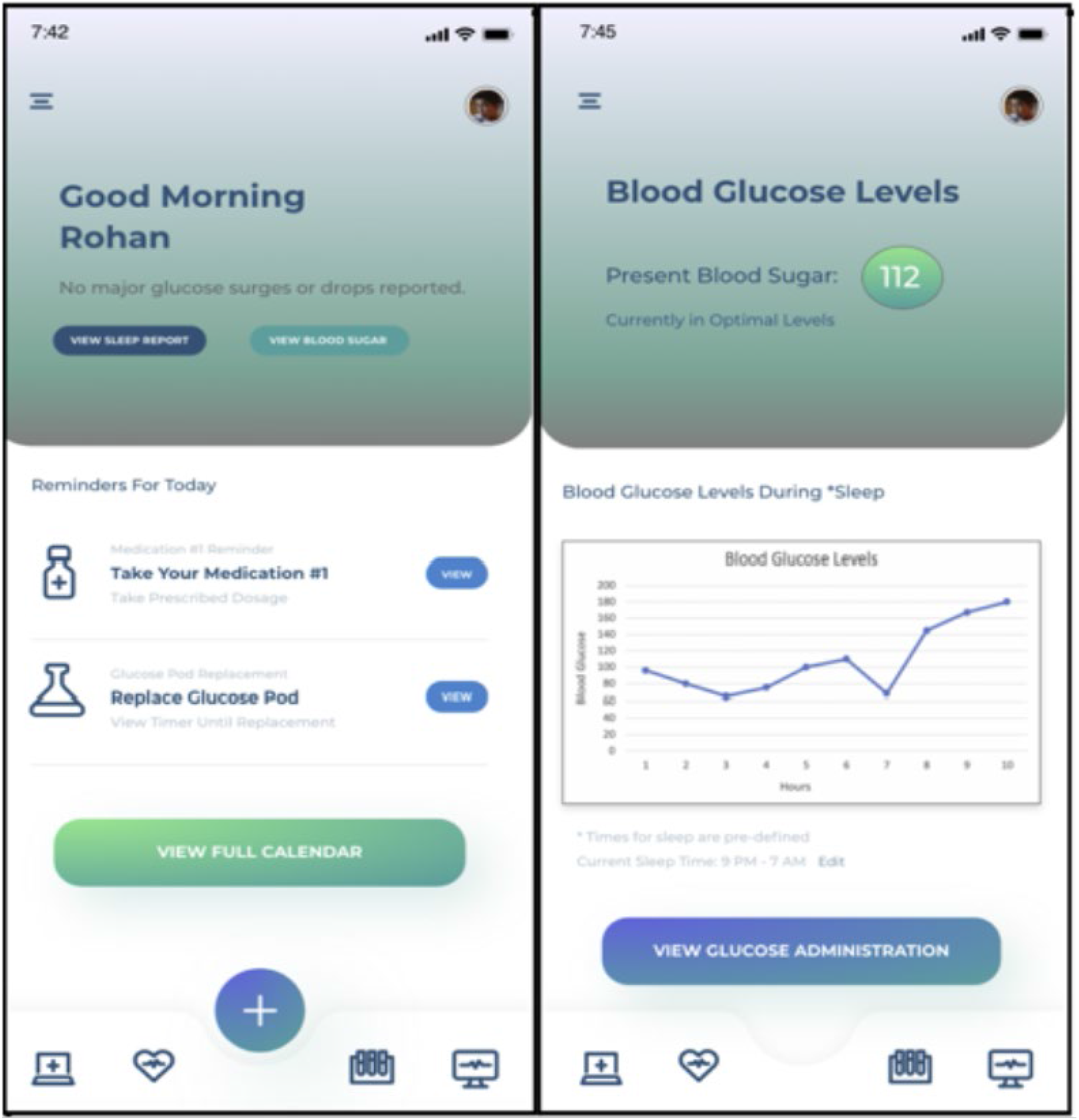
An example of the mobile app’s user interface developed in AdobeXD. The mobile app has the functionality to take in patient-specific data and blood glucose levels. Other features such as reminders and calendars are added for the user’s convenience.

### Deep Learning Model

Blood sugar, carbohydrate, and insulin bolus values are then sent to the deep learning model to predict future blood sugar levels. While blood sugar levels may appear to be spontaneous, levels usually correlate with lifestyle events (Reddy, P., H., 2017). For example, around mealtimes, patients with diabetes can expect a spike in blood sugar which is a primary reason why patients with diabetes often take insulin doses before or after meals (Donnor, T., et al., 2000). Blood sugar levels also may temporarily naturally fluctuate at certain times, rising or dropping and then returning to normal after a period (Mathew, TK., et al., 2023). Since glucagon and insulin do not have an immediate effect on the body, administering glucagon or insulin improperly during lifestyle events or natural fluctuations can lead to dangerous blood sugar levels (Slattery, D., et al., 2018; Ramnanan, C, J., et al., 2011). Treatment could have coincided with natural fluctuations in prior attempts utilizing glucagon and insulin in an artificial pancreas, resulting in continued cases of hyperglycemia and hypoglycemia (Haidar, A., et al., 2015; Haider, A., 2019; Jessica, R, Castle., et al., 2018; Lindkvist, E. B., et al, 2023).

Problems associated with the improper administration of insulin and glucagon can be avoided by administering treatment early using the model’s predictions. The deep learning model, built in Google Colab, utilizes the Python libraries TensorFlow (TensorFlow *v2*.*14*.*0*, 2023) and Keras (Keras documentation, 2023) for the model’s architecture while scikit-learn (*Documentation scikit-learn, 2023)* was used for data preprocessing and cleaning. The model is composed of two bidirectional Long Short-Term Memory (LSTM) network (BiLSTM) layers, one with 50 neurons and the other with a single neuron and utilizes Adam optimization. Compared to normal LSTMs and other model architectures, BiLSTMs allow for bidirectional information flow by processing input sequences not only from past to future but also from future to past. This allows the model to learn data patterns from both directions, enhancing the model’s understanding of temporal patterns (Xiao, D., et al., 2023). The BiLSTM layers culminate into a dense layer that provides us with the predicted values. The model is trained on data from the OhioT1DM dataset (Marling, C., & Bunescu, R., 2020) which contains 8 weeks’ worth of blood sugar, carbohydrate, and insulin data from 12 patients with type 1 diabetes. To further supplement the data available for the model, over 400,000 rows of synthetic blood sugar, carbohydrate, and insulin data were generated from the UVa/Padova Simulation (Man, C, D., et al., 2014). The model features dynamic training as real-time blood sugar, carbohydrate, and insulin dosage values are incorporated through continuous updates to optimize the model’s predictions. The model is trained with 400 epochs and a batch size of 32, demonstrating that it can be trained in a matter of minutes with the potential to be even faster with more powerful computing resources (Figure 6).

**Figure 6.**
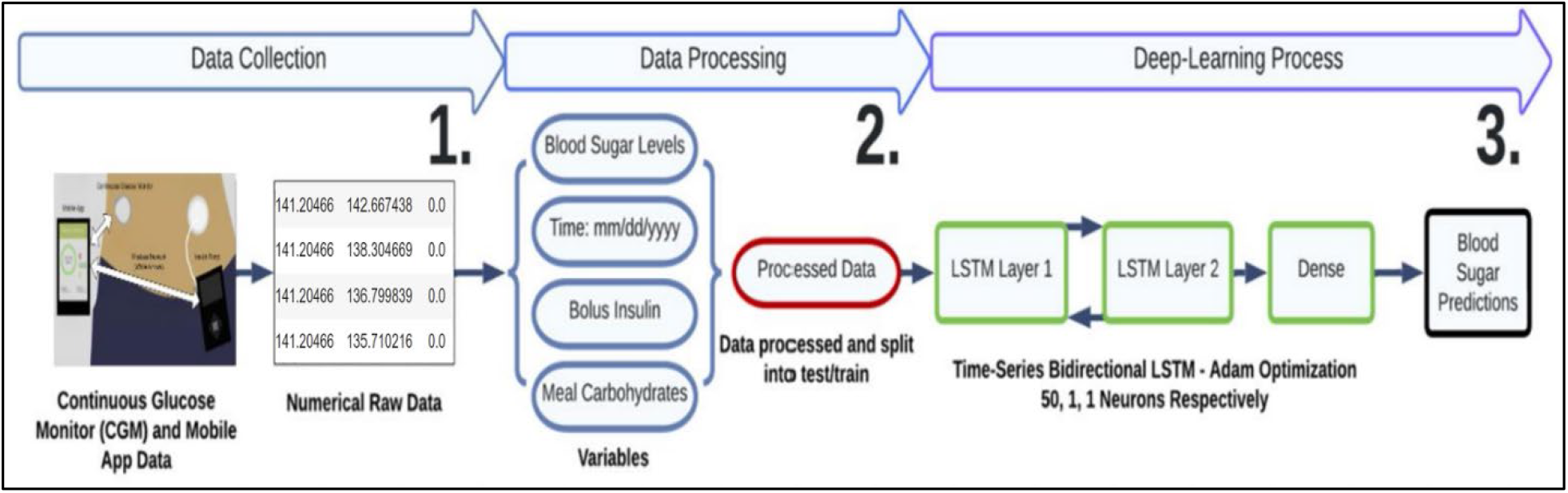
Flowchart of deep learning model’s three stages. First, (1) the insulin, carbohydrate, and blood sugar data are collected. The data is then processed (2) and fed into the time series deep learning model (3). The model outputs the final blood sugar predictions for the next 30 minutes.

### Dosage Algorithm

The deep learning model’s predictions are then sent to a Python algorithm to calculate the insulin and glucagon doses required. The algorithm starts by iterating through the predictions and determining if any of the values go above the maximum or below the minimum acceptable blood sugar ranges that the user had set in the mobile application. To account for natural or computational error, the slope of the values is calculated to determine if a user’s blood sugar is approaching a dangerous level. This helps prevent the improper administration of treatment during natural fluctuations as if the model’s predictions indicate that blood sugar will return to a normal level within 30 minutes, then treatment will not be delivered unless the patient’s blood sugar consistently remains elevated or low.

Using basic insulin and glucagon metabolization formulas along with a patient’s insulin and glucagon correction factors, which determine the effect of one unit of a hormone, dosage amounts are calculated and then sent via MQTT to the hardware insulin and glucagon pump to be administered (Koppanur, R., 2021; King, A. B., & Armstrong, D. U., 2007; Morris, CH., and Baker, J., 2023). Normally glucagon is delivered at a fixed rate, however, this system will deliver glucagon microdoses in a similar way that insulin is administered to allow for small increases in blood sugar. While the basic formulas provide an estimated dose calculation, more advanced mathematical functions will be needed in future algorithms to be more accurate (Figure 7).

**Figure 7.**
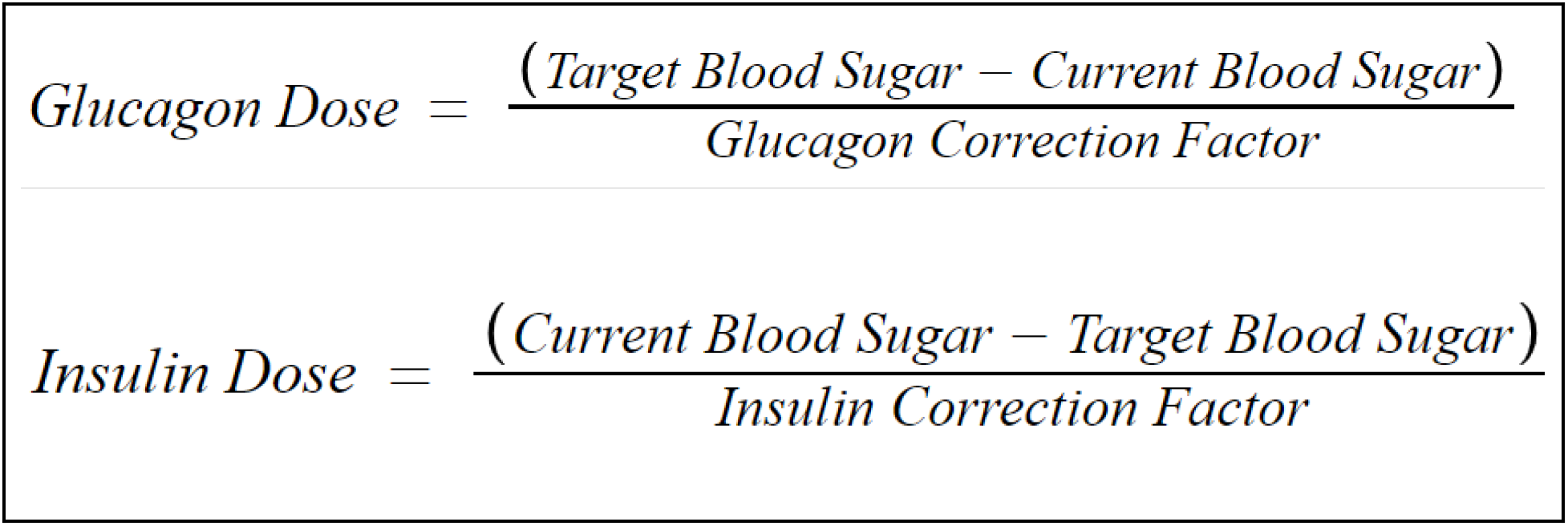
Basic insulin and glucagon dose calculation formulas. The glucagon and insulin doses take into consideration the target blood sugar and the current blood sugar doses. The correction factors in each of the equations are necessary to calculate the final dose. A fully deployed system would require more advanced mathematical functions (Koppanur, R., 2021).

### Secure MQTT Network

To defend against cyber threats and prevent data leakage, an MQ Telemetry Transport (MQTT) system was developed (Figure 8). MQTT is an internet protocol that allows the safe transfer of data between selected devices (Tsao, Y. C., 2022). The MQTT server was developed in Paho Python (Light, R., 2021) and is hosted using the HiveMQ Cloud Broker (HiveMQ documentation, 2023).

**Figure 8.**
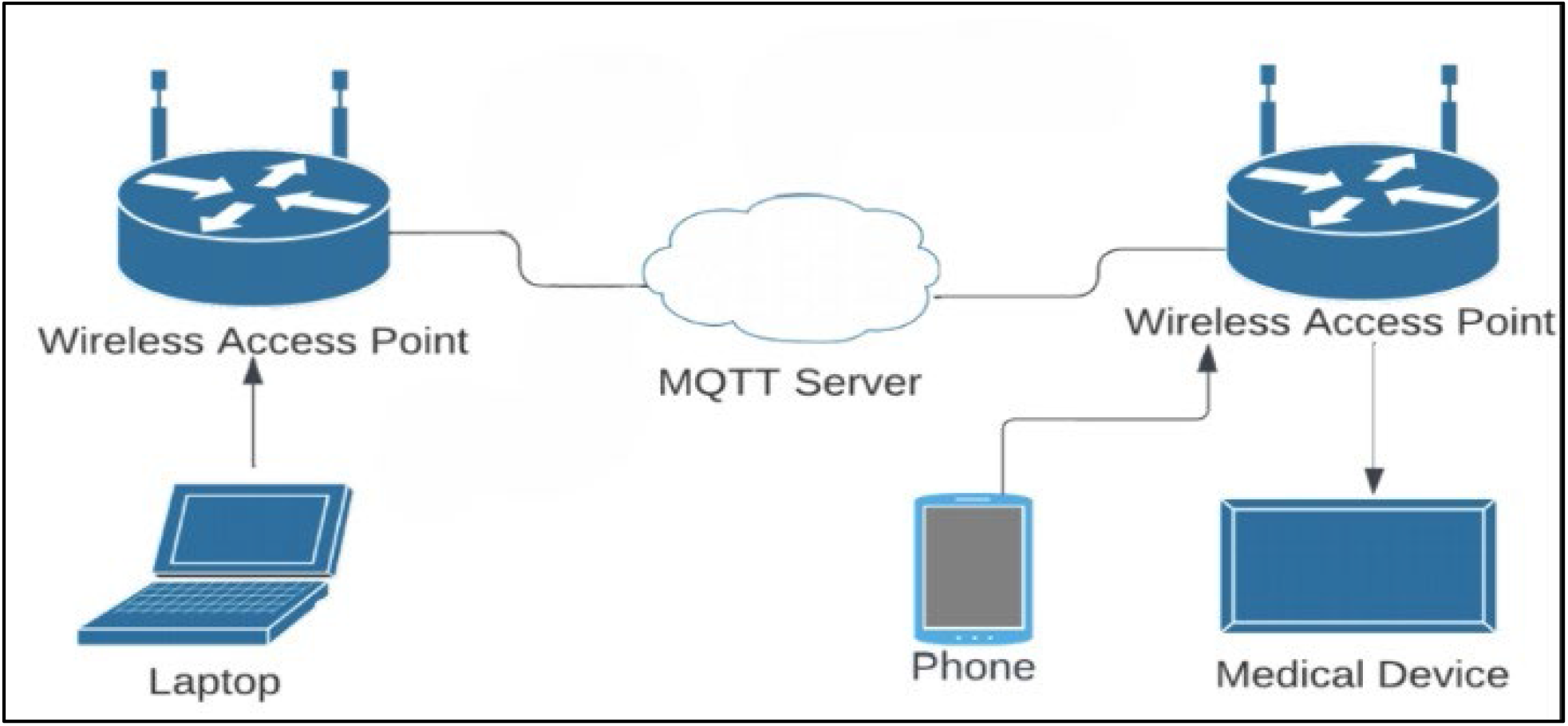
Diagram of the MQTT network. Devices on the network would need to be connected to the Internet so data can be transferred to and from the cloud-based MQTT server. The laptop refers to the system running the deep learning model and dosage algorithm, which would be a high-performance server.

The server has three layers of protection. First, all devices are hosted on the cloud and have their own unique ID and password. HiveMQ also employs numerous encryption protocols such as Transport Layer Security (TLS) which is a proven cryptographic protocol that allows for secure communication (Barker, E., and Barker, W, C., 2019). Between inter-device communications, the US National Security Agency and the National Institute of Standards and Technology (NIST)’s HMAC SHA256 algorithm is utilized to ensure the authenticity of devices on the network with unique signatures recognized by other devices. The HMAC SHA256 is currently unbroken and is used by world governments and cryptocurrency applications worldwide (A, L, Selvakumar., and C, S, Ganadhas., 2009). Each device has a unique HMAC SHA256 hash key, which cannot be reverted to its original form. This validates true devices on the network and ensures bad actors posing as normal system devices will not be able to receive or transfer data.

Serving as the final layer of defense, the Advanced Encryption Standard 256 (AES256) developed by NIST encrypts dosage, blood sugar data, and any other patient personal information to ensure patient safety and privacy. The AES256 is currently regarded as the industry standard of encryption and is used by world governments and militaries due to its current impenetrable nature (F, J, D’souza., and D, Pancha., 2017). All cybersecurity methods utilized on the MQTT server provide a comprehensive security system to protect the device from potential security breaches by malicious actors. While the system employed in this project was a prototype, real systems would utilize further methods such as key exchange systems for real-world deployment.

### Insulin and Glucagon Pump

The ultimate destination of the proposed system, the insulin and glucagon pump, is a standard insulin pump with added features to support a glucagon canister and tubing (Figure 9). To reduce costs, the shell and many of the basic components of the pump are made from biodegradable, extra-strong 3D-printer filament. Electronic components are soldered to microcontrollers without a breadboard to save space. The device is designed for the safe and effective delivery of insulin and glucagon using circuitry and mechanics. The device contains one Raspberry Pi Pico (Raspberry Pi Documentation, 2023) featuring WiFi connectivity allowing the device to receive data and execute movement and two mini-DC motors, powered by an L239D chip (L293D Data Sheet, 2023) which provides bidirectional current flow from the Raspberry Pi Pico, rotate at certain degrees to discharge the required amount of glucagon or insulin (Figure 10). The motors do this by driving a rack and pinion system attached to the canisters forward, similar to a plunger system, that releases glucagon or insulin. Even though the current prototype is fairly large, it can be miniaturized through printed circuit boards and smaller electronic components such as DC motors in the future.

**Figure 9.**
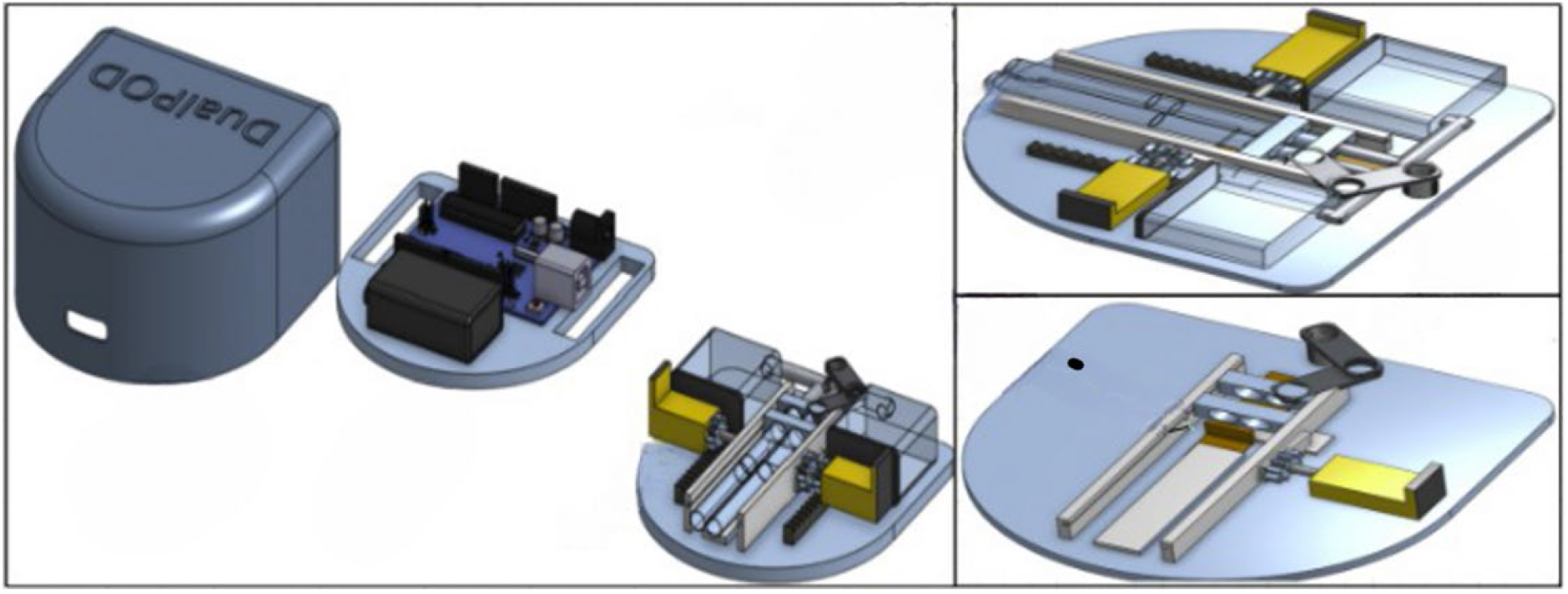
A general schematic of the insulin and glucagon pump. This Computer Aided Design (CAD) model was created using the Onshape software. The device features two levels in which the top level contains the device’s microcontroller and battery while the lower level contains the cannula and tube system in which glucagon and insulin get released.

**Figure 10.**
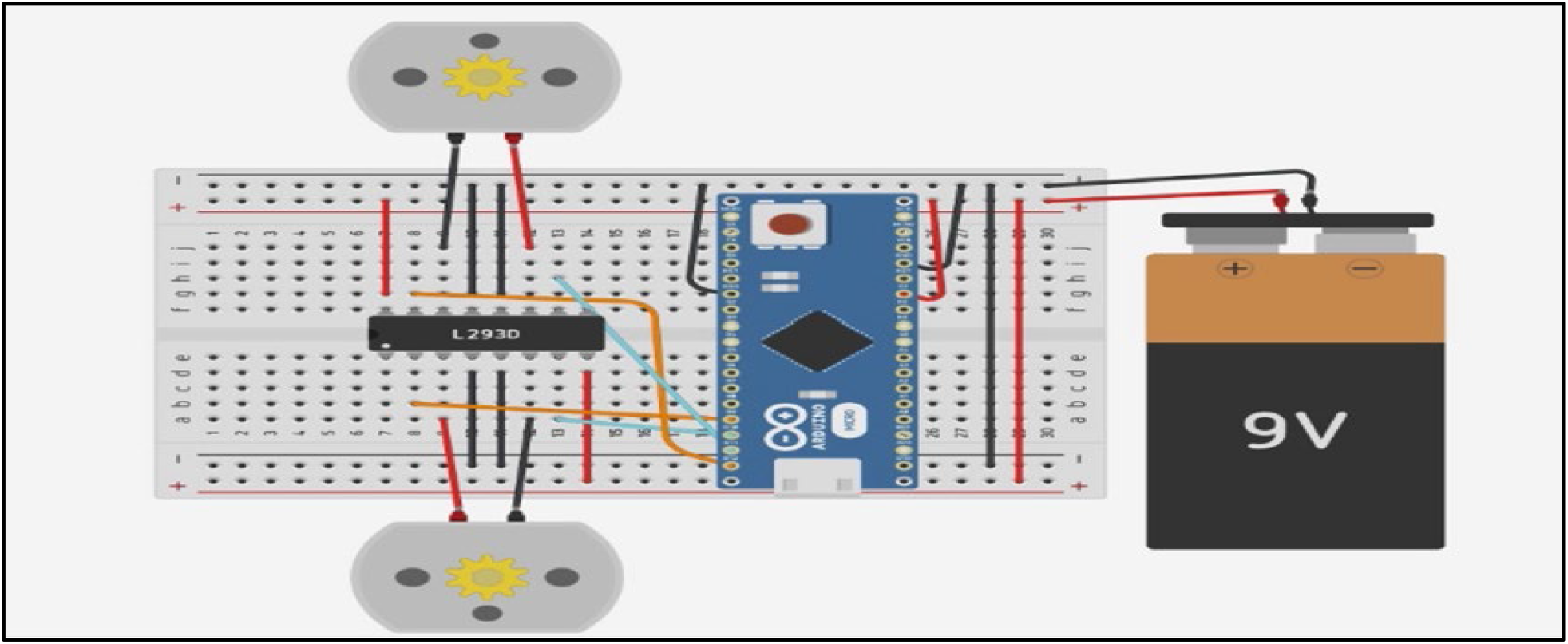
Insulin and Glucagon Pump Pinout diagram. The Arduino board depicted in the diagram represents where a microcontroller would be located. To power the two DC motors, an L293D chip is utilized to precisely distribute voltage to each motor. A 9-volt battery is also connected to the breadboard to power the microcontroller.

## Results

### Simulated Clinical Trials Overview and Results

The insulin and glucagon pump system was validated using the Python implementation of the FDA--approved preclinical trial alternative UVa/Padova Simulation (Man, C, D., et al., 2014). The system’s effectiveness was evaluated through a two-week randomized simulation involving 8 adults, 8 children, and 8 adolescents. Meals varied in both time and carbohydrate amount to test the system’s ability to dynamically respond to certain changes in lifestyle events. The trial was conducted again with a conventional Proportional-Integral-Derivative (PID) insulin pump being tested. PID works by analyzing deviations in blood sugar levels to calculate insulin doses and is one of the main types of control algorithms (Kang, S.L., et al., 2022).

As depicted in Figures 11 and 12, patients using the insulin-glucagon system compared to the patients using the PID system on average, spent (a) 38% more time in the normal blood sugar range, (b) 21.89% less time in the dangerous blood sugar range, (c) 9.56% less time in the low blood sugar range, and (d) 6.71% less time in the high blood sugar range. Since glucagon is not a feature in the model, only the deep learning model’s predictions were tested with insulin so future studies would need to modify the simulator to also evaluate glucagon and insulin.

**Figure 11.**
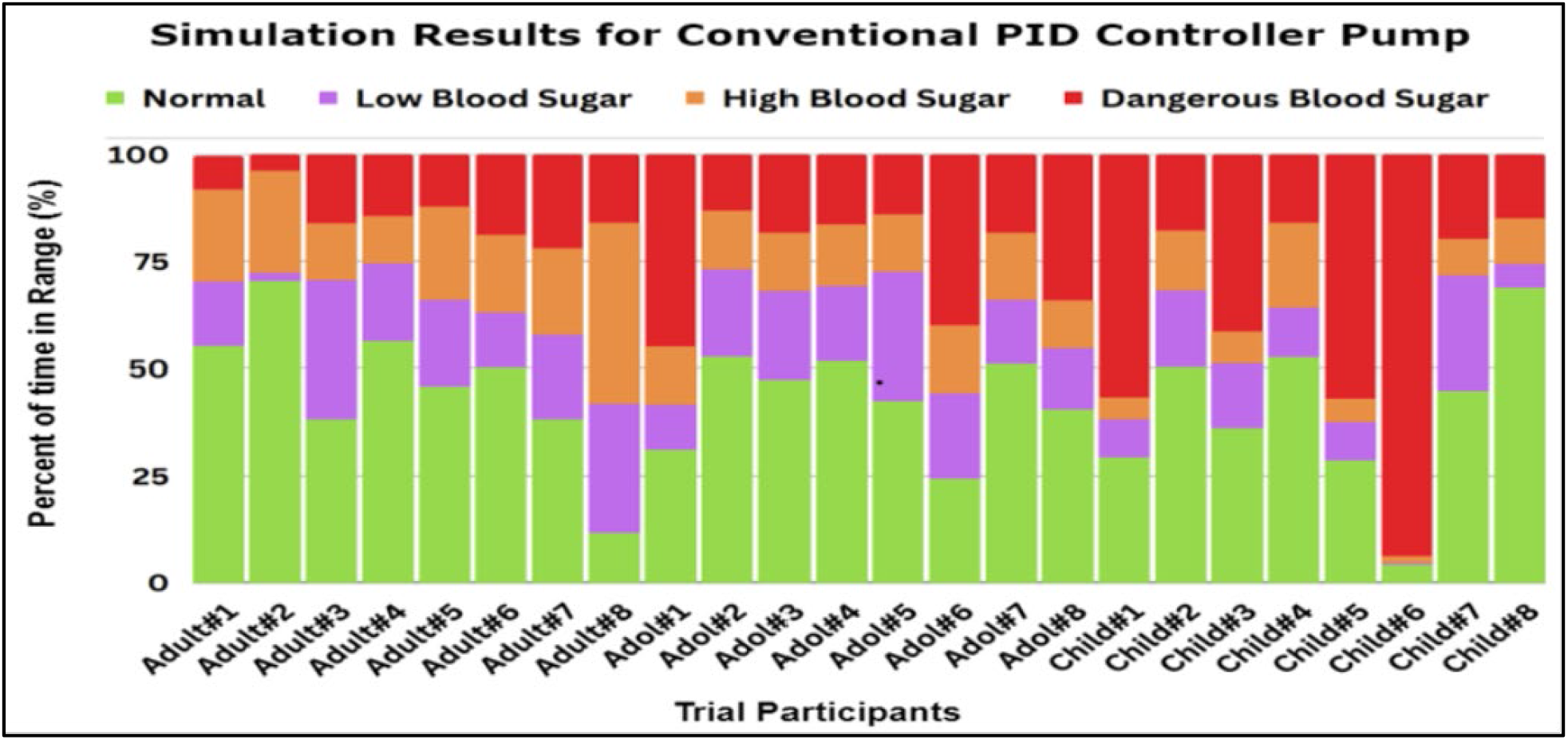
Results for the conventional PID controller pump’s effectiveness. The Python implementation of the Uva/Padova simulation contained a conventional PID controller which was tested. The graph shows that every in-silico patient tested spent time in the dangerous, low, and high blood sugar ranges.

**Figure 12.**
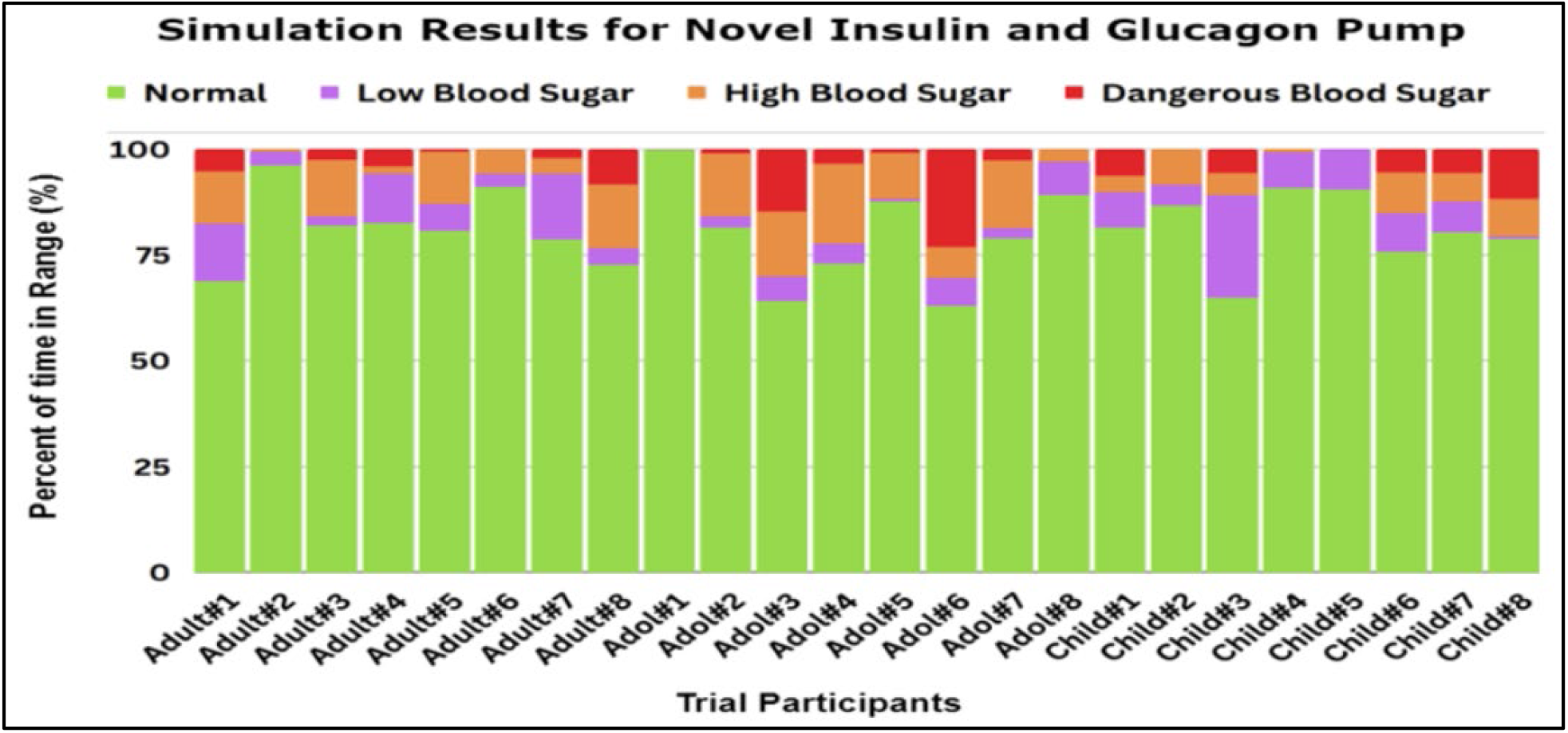
Results for the deep learning-based novel insulin and glucagon pump’s effectiveness. The deep learning model’s prediction aspect to better administer treatment was tested where compared to Figure 11, the in-silico trial participants were, on average, spending more time in the normal blood sugar range.

### Deep Learning Model Testing Results

To test the accuracy of the model, testing datasets containing new data from the OhioT1DM dataset and synthetic data from the UVa/Padova Simulation were utilized to evaluate accuracy. Utilizing both synthetic and real data allows the model to be evaluated in different contexts. Accuracy was evaluated in 30-minute intervals using Root-Mean-Squared-Error (RMSE) which was then converted into an accuracy percentage. The actual versus predicted values are depicted in Figure 13.

**Figure 13.**
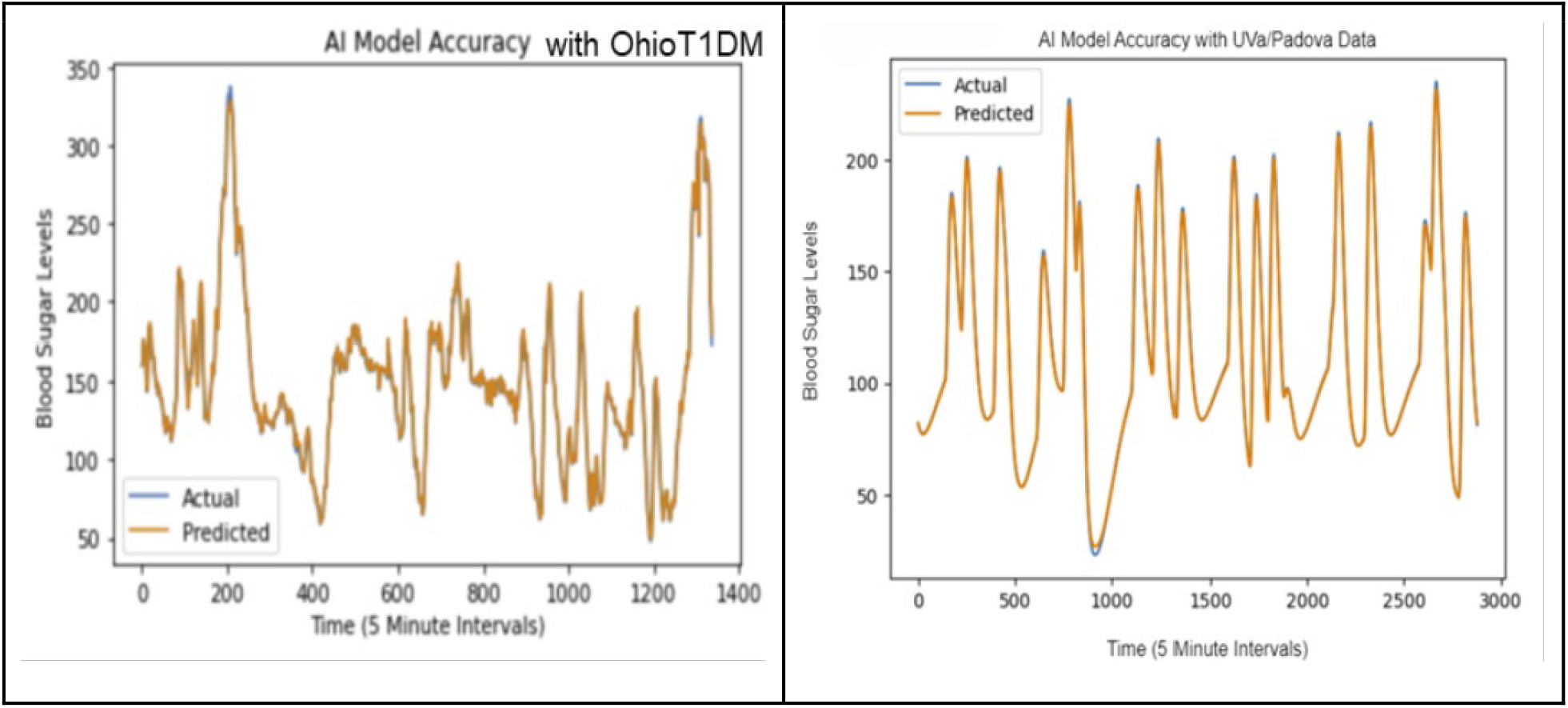
Deep learning model’s accuracy on both datasets. The graph on the left (a) represents the model’s accuracy with the OhioT1DM dataset and the graph on the right (b) represents the model’s accuracy with data from the Uva/Padova simulation. The orange line is the predicted value, and the blue line is the actual value. Lines in both graphs usually overlap, indicating the predictions are accurate in those instances.

The results from the training show that the highest achieved RMSE for the model (a) trained on OhioT1DM data in a single 30-minute interval: 1.233 mg/dl (∼96% Accuracy, and (b) trained on UVa/Padova data in a single 30-minute interval: 0.795 mg/dl (∼98°/o Accuracy). More advanced machine learning models will need to be developed to increase accuracy for a longer duration of time. The results show that the model was highly accurate on both datasets. The difference in accuracy could be attributed to the fact that the data from the OhioT1DM dataset is unprocessed, resulting in more noise and variability than the synthetic data from the UVa/Padova Simulation.

## Future Directions

In the future, certain aspects and components can be expanded, improved, and further tested to make this system safe and effective for patients with T1D. This includes improving the Deep Learning Model’s Accuracy through trying new model architectures such as Convolutional Neural Networks and different techniques such as transfer learning could be tested to develop more accurate blood sugar level prediction models. Increasing the accuracy and efficiency of these models would lead the system to administer proper doses and enact proper risk management. Creating an SQL database to save patients’ data would be beneficial for exploring future commercialization potential. However, this would also include new challenges in securing and managing this database. Future studies can modify the UVa/Padova simulation to test both insulin and glucagon. Further validating the medical device would be beneficial as while the medical device’s deep learning model and system were tested and validated in silica, future testing could involve human trials. In addition, the potential for commercialization of the device and approach will be explored.

## Conclusions

In this study, a novel artificial pancreas system has been developed and tested with a mobile application, deep learning model, dosage algorithm, secured MQTT network, and hardware prototype. Although it is reasonable to question the diverse range of technologies utilized in this project, it is in line with the rapid strides being made in personalized medicine approaches that take into consideration the differences in treatment strategies as per individual needs.

Having the ability to save lives and improve patient quality of life, this medical device holds enormous potential as a transformative tool for individuals with diabetes. It will enable patients to lead healthier and more fulfilling lives, free of the anxieties associated with diabetes management.

## Acknowledgments

The authors gratefully acknowledge access to the OhioT1DM dataset for testing the accuracy of the model. The data was obtained under a Data Use Agreement between the University of Arkansas and Ohio University for the purpose of conducting this research.

## Notes

### Competing Interest Statement

The authors have declared no competing interest.

